# Unraveling the dynamic transcriptomic changes during the dimorphic transition of *Talaromyces marneffei* through time-course analysis

**DOI:** 10.1101/2023.06.12.544618

**Authors:** Minghao Du, Changyu Tao, Xueyan Hu, Yun Zhang, Jun Kan, Juan Wang, Ence Yang

## Abstract

Systemic dimorphic fungi pose a significant public health challenge, causing over one million new infections annually. The dimorphic transition between saprophytic mycelia and pathogenic yeasts is strongly associated with the pathogenesis of dimorphic fungi. However, despite the dynamic nature of dimorphic transition, the current omics studies focused on dimorphic transition primarily employ static strategies, partly due to the lack of suitable dynamic analytical methods. Here, we firstly conducted time-course transcriptional profiling during the dimorphic transition of *Talaromyces marneffei*, a model organism for thermally dimorphic fungi. Then, we identified 5,223 dimorphic transition induced genes (DTIGs) by developing DyGAM-NS (dynamic optimized generalized additive model with natural cubic smoothing), a model that enables the capture of non-uniform and nonlinear transcriptional changes during intricate biological processes. Notably, the DyGAM-NS outperformed other commonly used models, achieving the highest F1-score in DTIGs identification. The cluster analysis of DTIGs suggests differential functional involvement of genes at distinct stages of dimorphic transition. Moreover, we observed divergent gene expression patterns between mycelium-to-yeast and yeast-to-mycelium transitions, indicating the asymmetrical nature of two transition directions. Additionally, leveraging the identified DTIGs, we constructed a regulatory network for the dimorphic transition and identified two zinc finger-containing transcription factors that potentially regulate dimorphic transition in *T. marneffei*. In summary, our study not only elucidates the dynamic changes in transcriptional profiles during the dimorphic transition of *T. marneffei* but also provides a novel perspective for unraveling the underlying mechanisms of fungal dimorphism.

**IMPORTANCE:** The dimorphic transition, i.e., morphological switch between saprophytic mycelia and pathogenic yeasts, plays a pivotal role in the pathogenesis of dimorphic fungi. However, the underlying mechanisms of dimorphic transition remain poorly understood, partly due to the lack of dynamic analytical methods suitable for its intricate nature. In the current study, we dissected the dynamic transcriptional profiles of dimorphic transition with a model thermally dimorphic fungus, *T. marneffei*, by developing a novel analytical method, DyGAM-NS. We proved that DyGAM-NS was more powerful in capturing the non-uniform and nonlinear gene expression variations during the dimorphic transition. With DyGAM-NS, we identified a repertoire of genes associated with dimorphic transition, and comprehensively unraveled distinct functions and expression patterns at different transition stages of *T. marneffei*, which offers novel perspectives regarding the mechanistic underpinnings of fungal dimorphism.

## Introduction

Systemic dimorphic fungi are significant human pathogens that cause millions of new infections annually worldwide (1). The ability of these fungi to transition between unicellular yeast and multicellular mycelial forms, known as dimorphic transition, enables them to adapt from environmental saprophytes to pathogens within hosts. In details, the mycelium-to-yeast (M-to-Y) transition is crucial for their pathogenicity as the yeast form evades the host immune system, while the yeast-to-mycelium (Y-to-M) transition is important for maintaining an environmental reservoir (2, 3). As the highly association between fungal dimorphic transition and their pathogenicity (3), elucidating the genetic mechanisms of dimorphic transition will not only shed light on understanding the pathogenic mechanisms of human fungal pathogens, but also provide theoretical guidance for molecular-based anti-fungal therapy.

The rapid advancement of high-throughput sequencing technologies, especially in the field of transcriptomics, has spurred numerous investigations aiming to employ more efficient genomic techniques for a comprehensive understanding of the regulatory mechanisms underlying fungal dimorphic transition (3). However, many studies have predominantly relied on static transcriptomic analysis approaches, focusing on contrasting transcriptomic differences between the mycelial and yeast phases to identify regulatory genes involved in the dimorphic transition process (4–7). Given the complex regulatory network governing fungal dimorphic transition, such static transcriptomic studies often fall short in capturing the full spectrum of genes that actively regulate the transition. While some studies have made efforts to elucidate the regulatory mechanisms by utilizing transcriptomic data obtained at different time points during the dimorphic transition, the limited number of sampling time points poses constraints on the comprehensive analysis of the intricate transcriptional changes that occur throughout the transition (8). Consequently, there is an urgent demand for innovative research strategies that can enable efficient and accurate deciphering of fungal dimorphic transition.

Temporal transcriptome analysis is a commonly employed approach for investigating dynamic gene expression changes in complex biological processes (9). It allows for the identification of transient gene expression patterns, response times, and gene regulatory relationships, thus holding significant potential for unraveling the regulatory mechanisms underlying dimorphic transition. However, the proper interpretation of time-course transcriptome data during dimorphic transition remains challenging. In recent years, various methods, such as polynomial regression (10), Gaussian process regression (11), autoregressive models (12), and natural cubic spline smoothing (13), have been developed to address the complexities associated with time- course transcriptome analysis. However, the dynamics of gene expression can differ greatly across species, particularly in microorganisms like fungi, where gene expression changes exhibit non-uniform and nonlinear patterns. This variability limits the effectiveness of existing models in capturing the complex temporal dependencies inherent in dimorphic transition. Therefore, it is essential to develop precise and adaptable methodologies that can effectively analyze time-course transcriptome data and uncover the dynamic gene expression changes occurring during dimorphic transition.

*Talaromyces marneffei* (formerly *Penicillium marneffei*) is a thermally dimorphic pathogenic fungus that undergoes a switch from a mycelial form at ambient temperature (25°C) to a yeast form at the host temperature (37°C) (14). Due to its lower biosafety level and the ease of inducing dimorphic transition in laboratory conditions, *T. marneffei* serves as an excellent model organism for studying fungal dimorphism (15). In this study, we employed a DyGAM-NS (dynamic optimized generalized additive model with natural cubic smoothing) approach to analyze the time-course transcriptome data during the dimorphic transition of *T. marneffei*. The DyGAM-NS model combines the flexibility of generalized additive models, which enable capturing nonlinear and complex changes, with dynamic optimization using natural cubic spline smoothing. By dynamically optimizing the model parameters for each gene, we identified the best-fitted models and determined the dimorphic transition induced genes (DTIGs) in *T. marneffei*. Cluster analysis of these DTIGs revealed dynamic expression patterns, providing insights into the temporal characteristics of the transcriptome during the dimorphic transition. Furthermore, by integrating the DTIGs with the transcription factors of *T. marneffei*, we constructed a gene expression regulatory network for the dimorphic transition, uncovering potential regulators involved in dimorphic transition of *T. marneffei*. In summary, the DyGAM-NS model not only enables efficient and accurate deciphering of the regulatory mechanisms underlying the dimorphic transition of *T. marneffei*, but also provides a powerful tool for unraveling complex biological processes in other microorganisms.

## Results

### Improving the reference genome of *Talaromyces marneffei* strain PM1 to near- chromosome level

To improve the quality of the reference genome for *Talaromyces marneffei* strain PM1 (8), we conducted high-coverage whole-genome sequencing using a combination of PacBio SMRT (3.3 Gb, 115×) and Illumina (6.0 Gb, 210×) sequencing technologies. Firstly, we employed PacBio data to construct the initial draft assembly, resulting in 26 contigs. Subsequently, we identified and eliminated 12 bubble sequences and one mitochondria sequence from the draft assembly. Finally, we polished the genome sequences using both PacBio raw data and high-coverage Illumina data. The final assembly consisted of 13 contiguous sequences, with a total size of 29.0 Mb, an N50 of 3.3 Mb, and an L95 of 8 (**Table 1**). Based on the Eurotiales database (*n* = 4,191), we successfully identified 4,067 complete BUSCOs (97.3%), indicating a significant improvement in the completeness and continuity of the PM1 genome sequences. Furthermore, we detected 16 telomere sequences (TCCTAA) and 8 centromere regions in the final assembly (**Figure 1**), consistent with previous findings (16, 17), thereby suggesting that the genome sequences have achieved near-chromosomal level.

**Figure 1.**
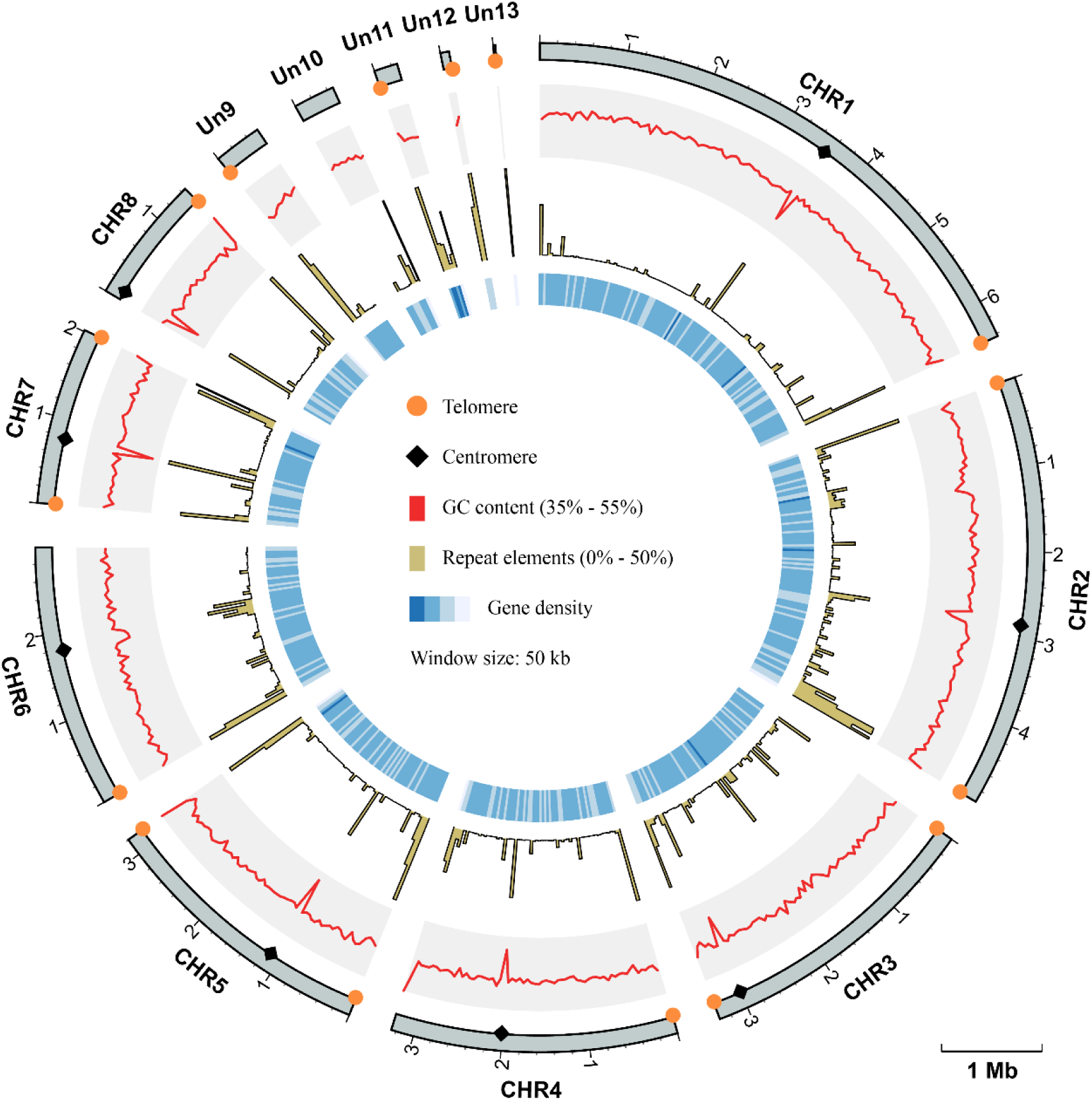
Circos plot of genome assembly and annotation of *T. marneffei* PM1. Each track, from outside to inside, represents: the coordinate of 13 sequences of the genome assembly of PM1, the GC contents (range from 35% to 55%), the proportion of repeat elements (range from 0 to 50%), and the gene density, respectively. All these genomic characteristics were analyzed using a window size of 50 kb. The telomere sequences are depicted by orange circles, while the centromere regions are represented by black diamonds.

**Table 1.**
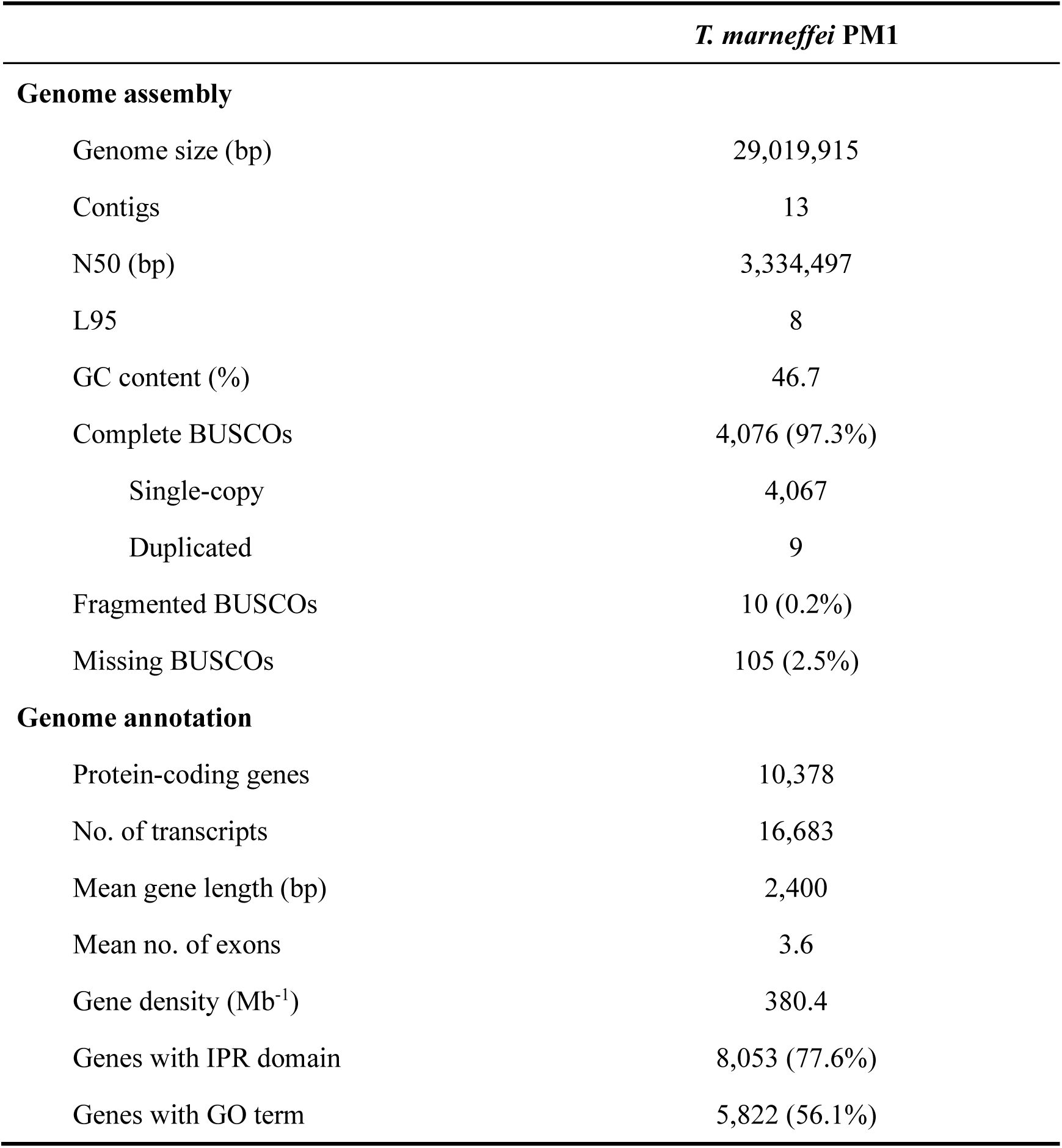
Summary of *T. marneffei* PM1 genome assembly and annotation

### Illustrating transcriptional dynamics of dimorphic transition by using time-course RNA-seq data

To explore the dynamic gene activity during the dimorphic transition of *T. marneffei*, we conducted RNA-seq analysis on samples collected at various stages of the time-course, encompassing both the mycelium-to-yeast (M-to-Y) and yeast-to-mycelium (Y-to-M) transitions. Recognizing the non-uniform nature of the dimorphic transition, we strategically selected 10 time points distributed unevenly over the 0−72 hours period (0, 1, 3, 6, 9, 12, 24, 36, 48, 72 hours) for both transitions. With three biological replicates for each time point, a total of 60 samples were subjected to RNA-seq, yielding an average of 24 million read pairs per sample.

Initially, these transcriptome data were employed to facilitate the genome annotation of PM1, resulting in the identification of 10,378 protein-coding genes (**Table 1**). Among these genes, 77.6% of the proteins were annotated with at least one InterPro domain, and 56.1% were assigned at least one Gene Ontology (GO) term. Subsequently, the RNA-seq data were mapped to the PM1 reference genome, and uniquely mapped read pairs were utilized to estimate the expression levels of protein-coding genes using the transcripts per million (TPM) method. To mitigate the impact of transcriptional noise arising from low-expression genes, we filtered out 9,541 genes with a mean TPM greater than one in at least one time point during either the M-to-Y or Y-to-M transition.

An initial analysis of the transcriptomic dynamics during the dimorphic transition was performed using principal component analysis (PCA). In the M-to-Y transition, PC1 accounted for 64.8% of the variance between samples. Arranging the samples in the order of sampling time revealed a distinct trajectory on the PC1-PC2 plane, indicating clear differentiation among the sample groups (**Figure 2A**). Notably, during the early stage of the M-to-Y transition (0−3 hours), the distances between sample groups were notably larger than those at other time points, suggesting a rapid transcriptional change during the initial phase of the M-to-Y transition. Similarly, in the Y-to-M transition, PC1 explained 61.1% of the variance between samples. A discernible trajectory was observed on the PC1-PC2 plane among the sample groups in the order of transition time (**Figure 2B**). The inter-group distances indicated that the magnitude of transcriptional changes during the early stage of the Y-to-M transition was not as pronounced as in the M-to-Y transition. Importantly, when all samples were subjected to PCA, a circular differentiation trajectory was formed between sample groups, aligning with the reversible cyclic nature of the dimorphic transition (**Figure 2C**). On the PC1-PC2 plane in the PCA of all samples, PC1 effectively distinguished between the M-to-Y and Y-to-M transitions, indicating distinct transcriptional regulation patterns during the dimorphic transition. Heatmap analysis based on gene expression levels across all samples further demonstrated that a considerable proportion of genes exhibited differential expression patterns during the dimorphic transition (**Figure 2D**), highlighting the complexity and nonlinearity of transcriptional dynamics during this process.

**Figure 2.**
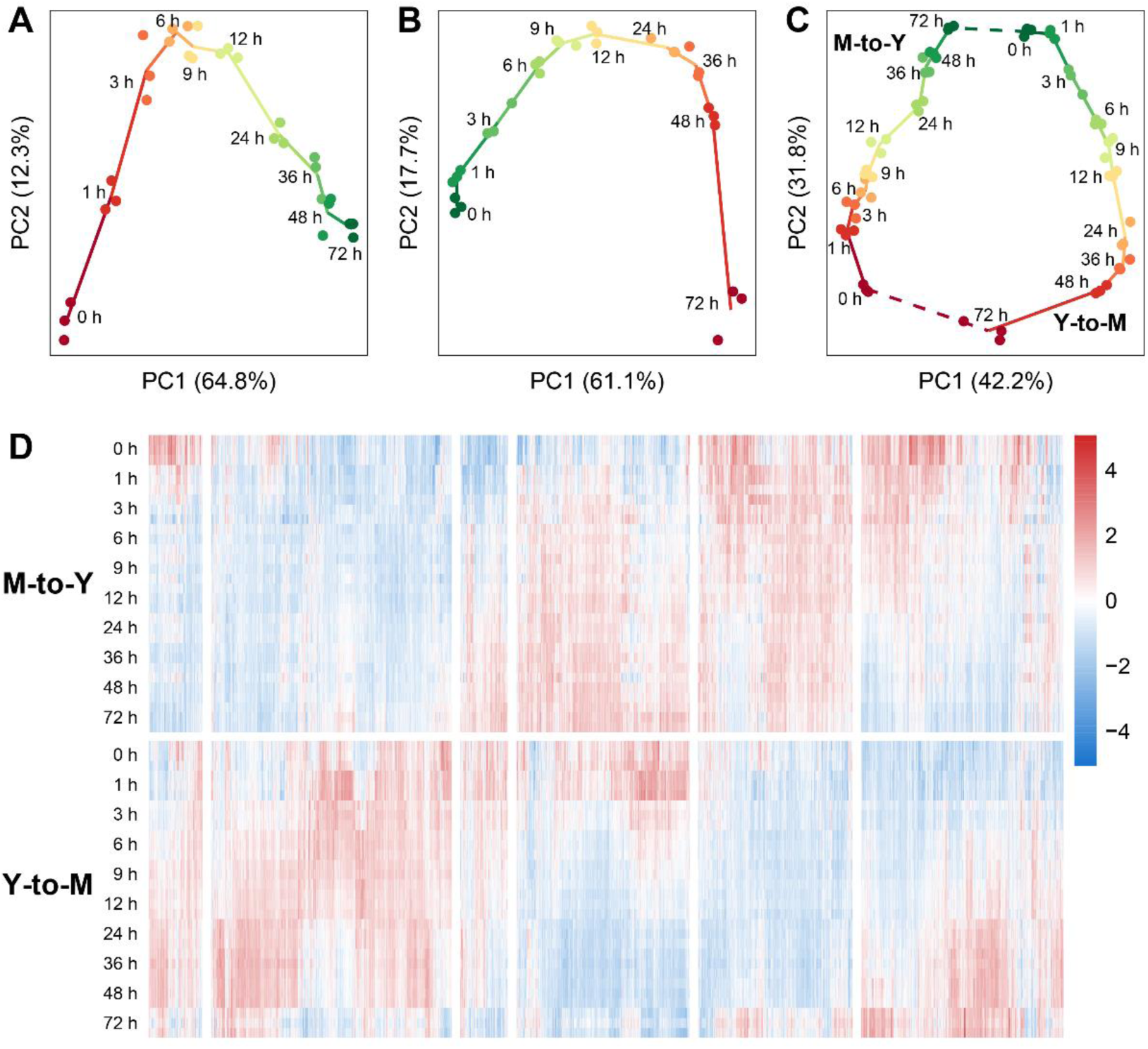
Transcriptional dynamics during M-to-Y and Y-to-M transitions of *T. marneffei*. The scatter charts show the differentiation trajectory during dimorphic transition of (A) M-to-Y, (B) Y-to-M, and (C) M-to-Y together with Y-to-M based on the scores of the first two principal components. (D) Heatmap of gene expression levels, measured with TPM, during M-to-Y and Y-to-M transition. The expression levels are standardized to have zero mean and unit variance across samples for each gene.

### Identification of dimorphic transition induced genes (DTIGs) by using generalized additive model and natural cubic spline

In this investigation, we employed an unevenly spaced sampling strategy that better aligns with the biological changes observed during the dimorphic transition of *T. marneffei*. To effectively capture the non-uniform and nonlinear dynamics of the time-course data, we proposed a dynamic fitting method, referred to as dynamic optimized generalized additive model with natural cubic smoothing (DyGAM-NS), to examine the dynamic alterations in gene expression induced by the dimorphic transition. By incorporating natural cubic splines as smoothing terms, DyGAM-NS allows for non-uniform and customizable partition knots, thereby accommodating the changing characteristics of gene expression during the dimorphic transition. The read counts for each gene during transition was modeled using a negative binomial distribution with gene-specific dispersion parameters. To explore the optimal parameters for the DyGAM-NS model, we randomly selected a subset of genes and fitted their expression changes over time using various NS partition numbers, NS knots, and dispersion coefficients. Model performance was evaluated using the corrected Akaike information criterion (AICc). The results demonstrated that gene-specific optimization of NS partition numbers, NS knots, and dispersion coefficients enhanced the performance of the DyGAM-NS model (**Figure S1**). Consequently, for each gene, we exhaustively explored combinations of NS smoothing term partition numbers and knots and selected the optimal parameter combination based on AICc. Compared to the default parameters, most genes (92.3% in M-to-Y and 96.4% in Y-to-M) exhibited improved model performance after parameter optimization (**Figure S2A and S2B**). Summarizing the optimal model parameters for each gene revealed that most of genes had an optimal partition number of 5 (**Figure S2C and S2D**), indicating the complex and diverse changes in the transcriptome during the dimorphic transition. In the M-to-Y transition, there was a noticeable presence of genes with partition knots at 1 and 3 hours compared to the Y-to-M transition (**Figure S2E and S2F**), further suggesting the more rapid early transcriptome changes in the M-to-Y transition.

To identify genes specifically induced by the dimorphic transition (DTIGs), we applied stringent filtering criteria to the DyGAM-NS model and identified 3,084 and 4,156 DTIGs in the M-to-Y and Y-to-M transitions, respectively. Due to the lack of known data on true DTIGs of *T. marneffei*, direct evaluation of the precision and recall of the DyGAM-NS model is challenging. Thus, we alternatively estimated precision and recall using two different approaches. Given that genes involved in regulating the dimorphic transition should display significant expression changes during the process, DTIGs that do not exhibit differential expression between any two time points in the time-course data are likely false positives. Applying this criterion, we estimated the precision of M-to-Y DTIGs to be 99.7% (3,076/3,084) and Y-to-M DTIGs to be 99.8% (4,146/4,156) (**Table 2**). To assess the recall of the model, we identified highly variable genes (HVGs) based on the biological coefficient of variation (BCV) during the dimorphic transition. Genes involved in regulating the dimorphic transition are expected to have higher BCV in temporal data compared to a static state. We employed a chi-square test to identify genes with significantly higher (FDR < 0.05) BCV (18) during the dimorphic transition than in a static state, considering them as HVGs. Using the HVG dataset as the criterion to evaluate the DyGAM-NS model, we estimated the recall of M-to-Y to be 90.3% (140/155) and Y-to-M to be 91.6% (261/285) (**Table 2**). Finally, based on the estimated precision and recall, the F1-scores of the DyGAM-NS model in the M-to-Y and Y-to-M transitions were 94.7% and 95.4%, respectively (**Table 2**).

**Table 2.**
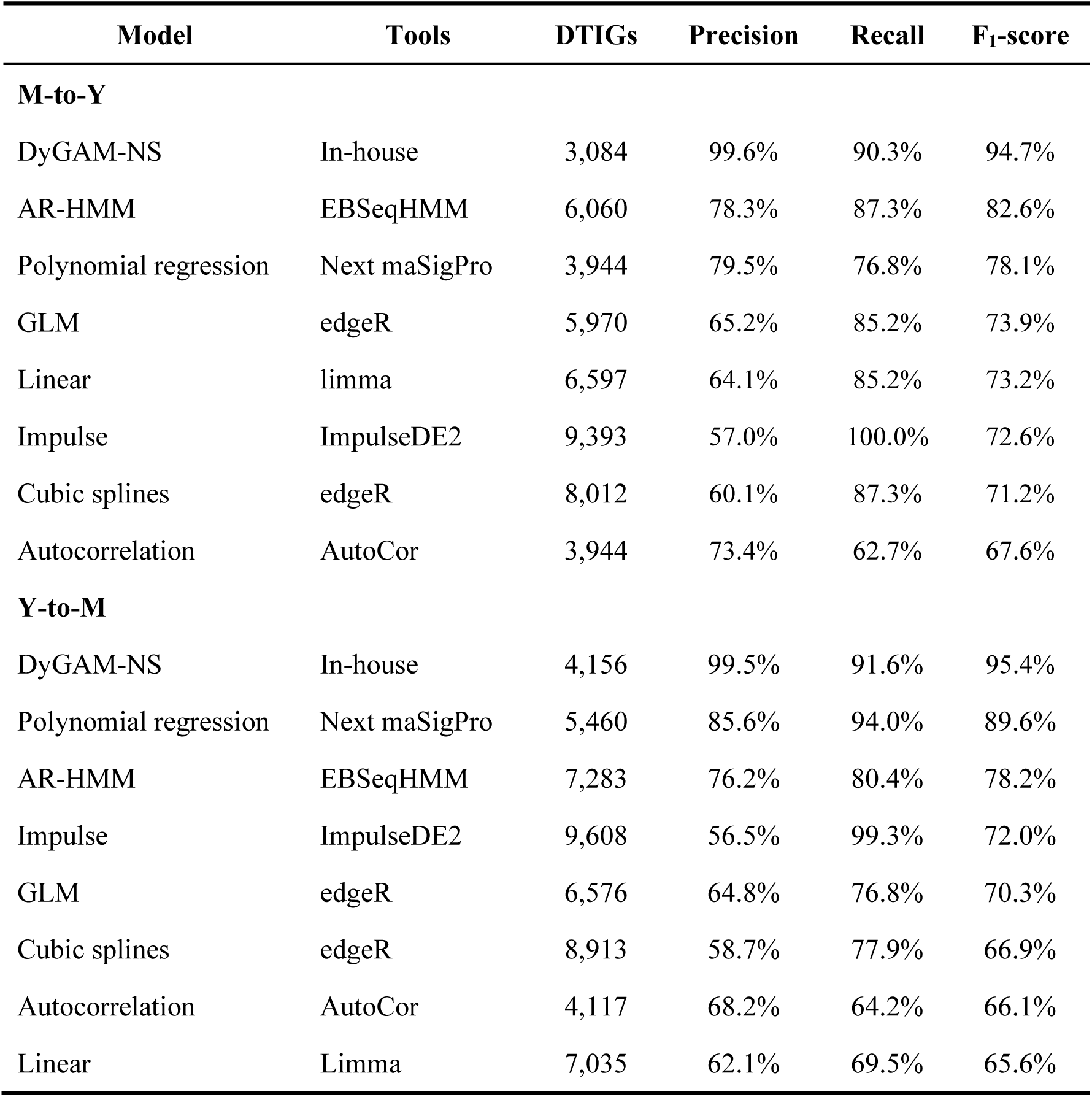
Performance of DyGAM-NS and other commonly used models in

To comprehensively assess the effectiveness of the DyGAM-NS model in identifying DTIGs, we conducted a comparison with seven commonly employed time-course analysis methods (10, 12, 19–21) (**Table 2**). The findings revealed that the DyGAM-NS model exhibited the highest precision in both the M-to-Y and Y-to-M transitions. Regarding recall, the DyGAM-NS model also demonstrated favorable performance. While the ImpulseDE2 model displayed higher recall than the DyGAM- NS model, its lower precision led to suboptimal overall performance. Overall, the DyGAM-NS model achieved the highest F1-score in both the M-to-Y and Y-to-M transitions, indicating its efficacy as a DTIGs identification model. Furthermore, when comparing the different models used to identify DTIGs (**Figure S3**), it was observed that a large majority of DTIGs (99.1% in M-to-Y and 98.8% in Y-to-M) identified by the DyGAM-NS model were also identified as DTIGs by at least one other method, highlighting the robust performance of DyGAM-NS model.

### Clustering of DTIGs indicate that different gene expression pattern for M-to-Y and Y-to-M transitions of *T. marneffei*

To explore the dynamic changes of DTIGs during the dimorphic transition of *T. marneffei*, we employed a modified trimmed *k*-means clustering algorithm (22) to examine their expression patterns over time. In the M-to-Y transition, we discovered 11 distinct expression patterns (**Figure 3**), encompassing three primary types of changes: up-regulated (m2y_P2, m2y_P5, and m2y_P8), down-regulated (m2y_P1, m2y_P3, m2y_P6, m2y_P7, and m2y_P9), and pulse-like (m2y_P10 and m2y_P11). Among the up-regulated expression patterns, we observed critical genes involved in the survival of *T. marneffei* within host macrophages, such as *pepA* (23) in pattern m2y_P5, iron affinity-related genes (24) (*ftrA*, *ftrC*, and *sidA*) in pattern m2y_P2, cytochrome P450 monoxygenases (25) (*simA*), and the tyrosine metabolic regulator (26) (*hmgR*) in pattern m2y_P8. These findings suggest significant transcriptional changes occurring during the early stages of the M-to-Y transition, facilitating rapid adaptation to the harsh intracellular environment. Concerning the down-regulated expression patterns, we identified genes associated with conidiation and pigment synthesis (27–31) (*alb1*, *pbrB*, *pks11*, *pks12*, *brlA*, and *stuA*), which are closely linked to the morphological characteristics of the mycelia. Additionally, the pulse-like expression patterns exhibited an immediate transient decrease in gene expression at the early stage of the M-to-Y transition, potentially indicating a transient response regulation mechanism during this transition. However, the underlying mechanism governing these genes remains unknown and warrants further investigation.

**Figure 3.**
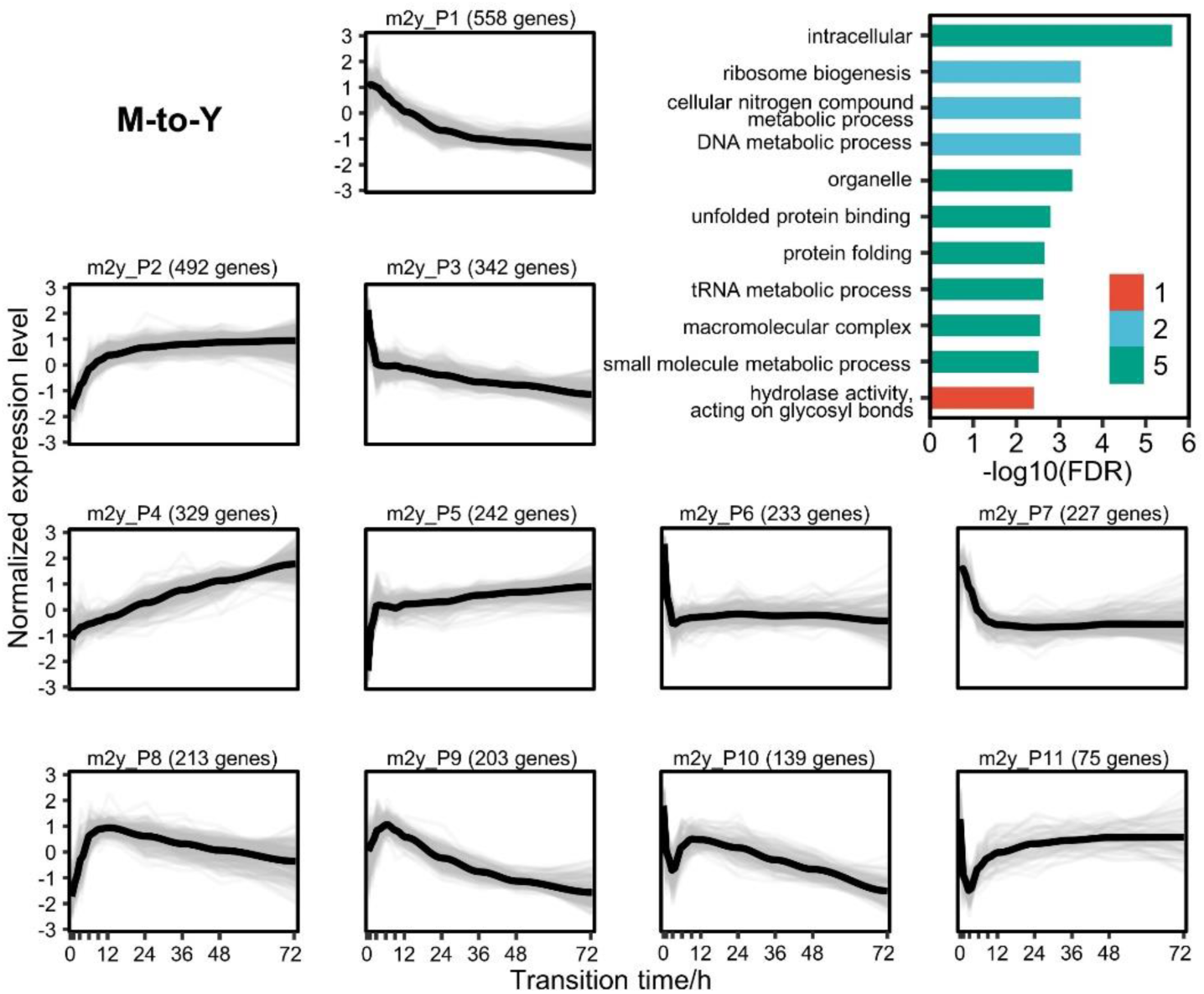
Gene expression levels and enriched gene ontologies of the 11 expression patterns in M-to-Y transition. The line charts represent the expression tendency of the 11 expression patterns in M-to-Y transition. Each light grey line represents a gene, and the black lines are drawn based on the mean values at each time point. The gene expressions levels are measured with TPM and standardized to have zero mean and unit variance across samples. The bar chart shows enriched gene ontologies for each expression pattern, which are presented in different colors.

DTIGs associated with the Y-to-M transition can be classified into 11 distinct expression patterns (**Figure 4**), which can further be grouped into three types of changes: up-regulated (y2m_P2, y2m_P3, y2m_P4, y2m_P5, and y2m_P8), down-regulated (y2m_P1 and y2m_P6), and pulse-like (y2m_P7, y2m_P9, y2m_P10, and y2m_P11). In the up-regulated expression pattern y2m_P2, we identified two basic leucine-zipper (bZIP) transcription factors (32, 33) (*atfA* and *yapA*), known to regulate the response of *T. marneffei* to environmental stress. Another up-regulated pattern, y2m_P4, contains the C2H2 transcription factor (34) (*hgrA*), which governs the formation of the hyphal cell wall. Similarly, up-regulated patterns y2m_P3 (29, 31, 35) (*gasC*, *pks11*, *pks12*, and *stuA*) and y2m_P5 (27, 28, 30) (*pbrB*, *alb1*, and *brlA*) exhibit genes related to the morphological characteristics of the mycelia, including conidiation and pigment synthesis. Notably, the dynamic changes in the transcriptome during the Y-to-M transition are not as pronounced as those observed in the M-to-Y transition. For instance, the mycelia phase-related genes found in the y2m_P5 expression pattern gradually up- regulate in the later stage of the Y-to-M transition (**Figure 4**). In terms of the down- regulated expression patterns, functional enrichment analysis reveals a significant enrichment of translation-related functions, indicating a decreased demand for protein synthesis and related processes during the Y-to-M transition. Moreover, the continuously down-regulated pattern y2m_P6 exhibits a significant enrichment of mitochondrial-related components and genes associated with the tricarboxylic acid (TCA) cycle. This finding is consistent with the fact that *T. marneffei* relies primarily on the TCA cycle for energy metabolism in the yeast phase, whereas glycolysis is prominent in the mycelia phase (36). Additionally, we observed genes related to intracellular survival (23, 24) (*sidA*, *pepA*, and *ftrC*) in the two down-regulated expression patterns. Within the pulse-like expression patterns, y2m_P7 includes the RFX transcription factor (37) *rfxA*, which plays a role in regulating cell division. Functional enrichment analysis also indicates a significant enrichment of cell division and cycle-related functions in y2m_P7, suggesting that cell division is inhibited in the early stage of the Y-to-M transition and reactivated in the later stage.

**Figure 4.**
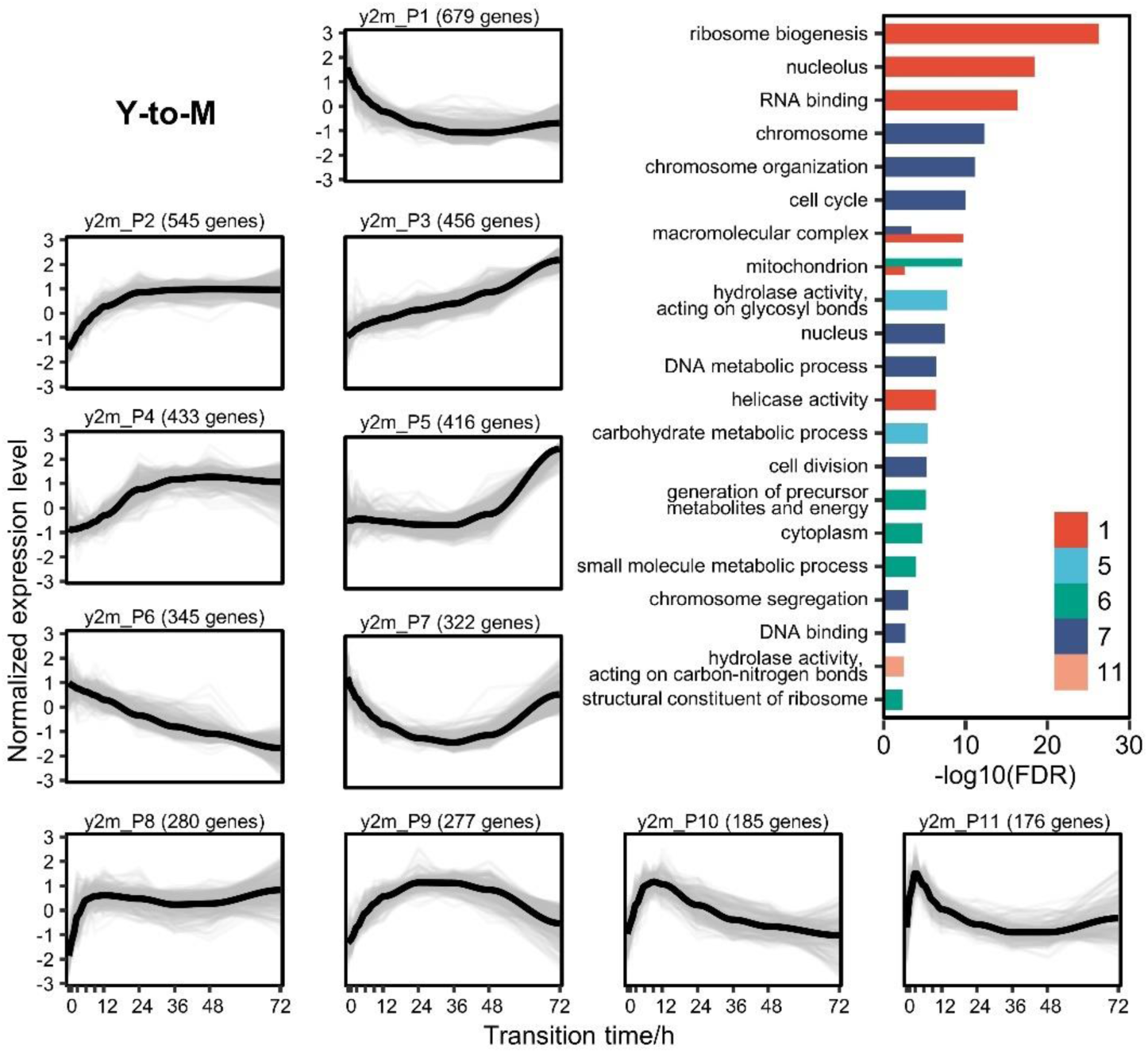
Gene expression levels and enriched gene ontologies of the 11 expression patterns in Y-to-M transition. The line charts represent the expression tendency of the 11 expression patterns in Y-to-M transition. Each light grey line represents a gene, and the black lines are drawn based on the mean values at each time point. The gene expressions levels are measured with TPM and standardized to have zero mean and unit variance across samples. The bar chart shows enriched gene ontologies for each expression pattern, which are presented in different colors.

### Gene regulation network underlying potential regulators of dimorphic transition of *T. marneffei*

To elucidate the regulatory relationships among DTIGs during the dimorphic transition, we employed the GRNboost2 (38) method to construct a directed gene regulatory network. To enhance the accuracy of the GRNboost2 approach, we utilized 5,223 DTIGs identified by the DyGAM-NS model in both the M-to-Y and Y-to-M transitions as target genes. Additionally, we incorporated 401 candidate transcription factors as prior information, identified through a combination of functional annotation of *T. marneffei* and known fungal transcription factor domains (39). To ensure the focus on robust regulatory relationships, we retained the top 5% cumulative weight of strong gene-gene regulatory interactions to form the final gene regulatory network. Following the filtering process, the dimorphic transition gene regulatory network comprised 2,243 pairs of robust regulatory relationships involving 1,664 genes, of which 325 were transcription factors (**Figure 5A**). Notably, within these 1,664 genes, TM060272, TM021494, and TM030196 emerged as the most significant genes based on their betweenness centrality within the network (**Figure 5B**). TM060272, possessing a Zn(2)-C6 fungal-type DNA-binding domain (IPR001138), displayed up-regulation during the M-to-Y transition but exhibited minimal changes during the Y-to-M transition, indicating its role in M-to-Y transition regulation. Conversely, TM021494, which also contains a zinc finger domain, exhibited up-regulation during the Y-to-M transition while showing little change during the M-to-Y transition, suggesting its involvement in the regulation of the Y-to-M transition. The TM030196 gene (*abaA*) exhibited an immediate transient increase in expression during the early stage of the M- to-Y transition (0−9 hours) but showed minimal changes during the Y-to-M transition, supporting its regulatory role in the M-to-Y transition, consistent with previous research (40).

**Figure 5.**
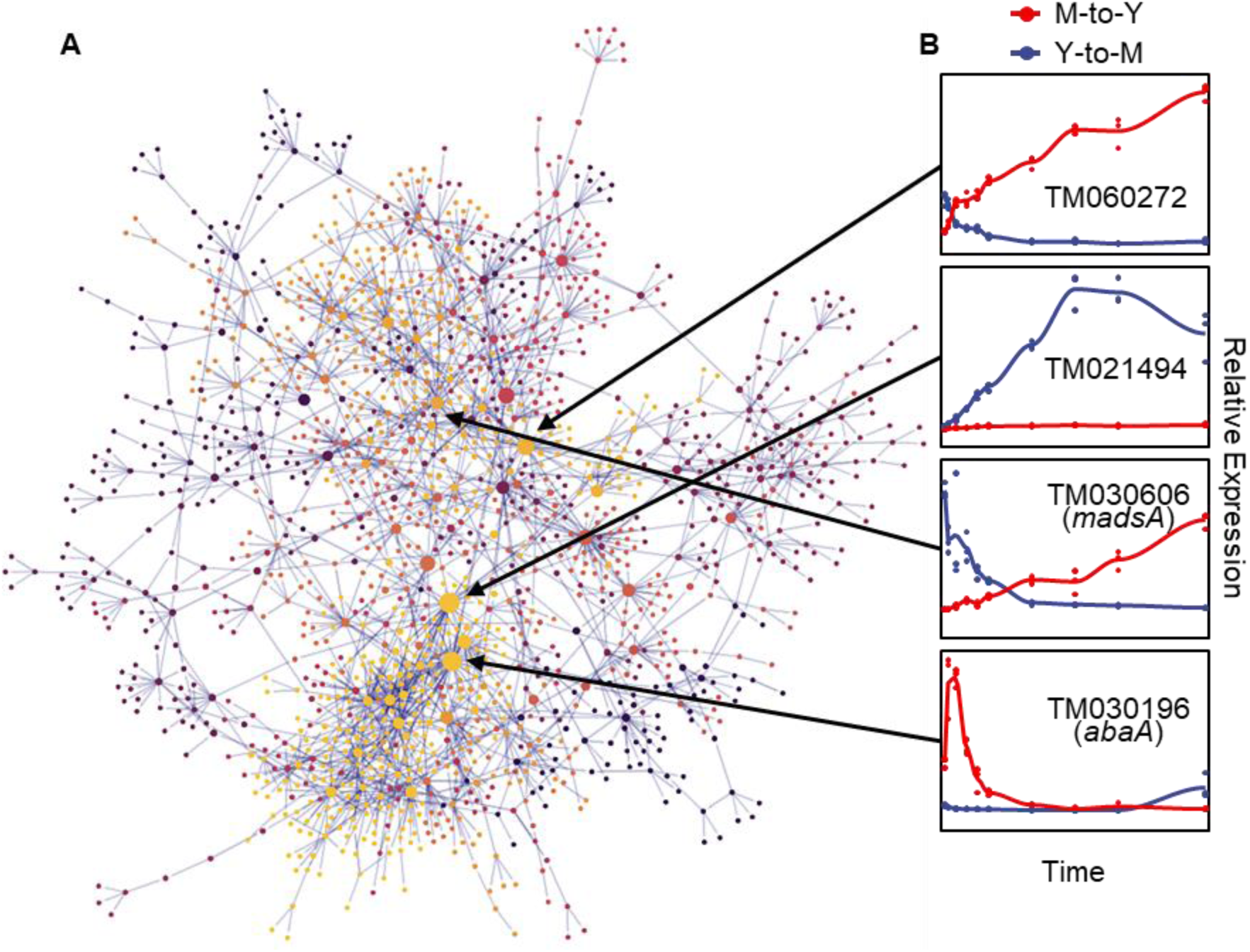
Gene regulatory network of dimorphic transition of *T. marneffei*. (A) Visualization of gene regulatory network of dimorphic transition. Each point represents a gene, and the size of points represents importance of genes in the network, which is measured with betweenness. The light grey lines represent strong regulatory relationship between corresponding gene pairs. (B) The line charts on the right part represent tendency of relative expression level (measured with TPM and standardized across samples) in M-to-Y (red line) and Y-to-M (blue line) transition for four putative regulators: TM060272, TM021494, TM030606 (*madsA*), and TM030196 (*abaA*).

To assess the precision of the gene regulatory network governing the dimorphic transition, we focused on the MADS-box transcription factor *madsA* gene (TM030606), which has been experimentally verified to play a role in regulating the dimorphic transition (8, 41) (**Figure 5B**). To validate the downstream regulatory genes within the dimorphic transition regulatory network, we employed knockout and overexpression strains of *madsA*. RNA-seq analysis was conducted on these strains at four time points during the dimorphic transition (0 and 48 hours for M-to-Y, and 0 and 6 hours for Y-to- M). By integrating the transcriptome data from the wild-type strain at corresponding time points, we identified 1,788 genes regulated by *madsA* during the dimorphic transition, of which 1,066 were identified as DTIGs by DyGAM-NS. Notably, in the gene regulatory network constructed using GRNboost2, over 50% (7/12) of the downstream target genes of *madsA* were validated (**Figure S4**).

## Discussion

Dimorphic transition represents a sophisticated adaptive mechanism in thermally dimorphic fungi, enabling their survival and propagation across diverse environments. Understanding the regulatory mechanisms underlying fungal dimorphic transition is a key scientific pursuit, given its close connection to fungal pathogenicity. Our study focused on the model organism of dimorphic fungi, *Talaromyces marneffei*, and demonstrated the significant potential of high-resolution temporal transcriptomes in deciphering the regulatory mechanisms underlying fungal dimorphic transition. On the one hand, temporal transcriptomes provide comprehensive coverage of the entire biological process, surpassing the limitations of static transcriptomic studies. In previous studies, comparisons of transcriptional profiles at different temperatures yielded limited information, with only a fraction of genes associated with dimorphic transition identified (7, 8). In contrast, our study utilized temporal transcriptional data spanning the first 72 hours of M-to-Y and Y-to-M transitions, resulting in the identification of a larger number of dimorphic transition induced genes (DTIGs). Specifically, we identified 3,084 DTIGs in the M-to-Y transition and 4,156 DTIGs in the Y-to-M transition, accounting for 50.3% (5,223 out of 10,378) of all protein-coding genes. This substantial increase in DTIG identification highlights the importance of high-resolution temporal data in comprehensively identifying regulatory genes related to dimorphic transition. On the other hand, temporal data also enables the observation of gene expression changes throughout the biological process, facilitating the discovery of distinct expression patterns during dimorphic transition. Our differentiation trajectory analysis revealed divergent gene expression patterns between the M-to-Y and Y-to-M transitions. Moreover, we identified numerous transition-specific DTIGs (1,067 for M-to-Y and 2,139 for Y-to-M), further supporting the asymmetrical nature of the two transition directions. Furthermore, the clustering of DTIGs in both the transition from M-to-Y and Y-to-M reveals the presence of numerous distinctive expression patterns throughout the dimorphic transition, including rapid up or down-regulation, continuous up or down-regulation, and pulse-like. Through functional analysis, it becomes evident that diverse gene functions are significantly enriched within these distinct expression patterns, suggesting that different biological functions play prominent roles at different stages of dimorphic transition. Consequently, our study underscores the criticality of temporal transcriptomic data as an indispensable tool for dissecting the intricate nature of fungal dimorphic transitions.

Despite the decreasing cost of sequencing technologies, which enables high-resolution time-course transcriptome sequencing, there remain challenges in deciphering temporal data for complex biological processes. In this study, we developed the DyGAM-NS model, which combines a flexible generalized additive model and dynamic gene-specific natural cubic spline fitting optimization. We compared the DyGAM-NS model with seven other commonly used time-course analysis methods. To evaluate the performance of the DyGAM-NS model, we utilized differentially expressed gene sets identified through pairwise comparisons at each time point and highly variable gene sets based on biological coefficient of variation (18) as benchmarks, since a standardized dataset for DTIGs in *T. marneffei* was not available. Although ImpulseDE2, based on impulse model had the highest recall rate, it also had a high false positive rate, resulting in an accuracy of less than 60% for the model. Previous studies have also indicated that the false positive rate of ImpulseDE2 increases with the number of sampling time points (42). This phenomenon may be attributed to the inadequacy of the pulse model in fitting complex temporal data, leading to low accuracy in identifying DTIGs. The spline smoothing fitting is theoretically more suitable for analyzing temporal data with complex variations. However, most current spline-based time-course analysis methods adopt uniform partition fitting and lack optimization of partition numbers, resulting in poor performance in analyzing the temporal data of dimorphic transition. In contrast, our proposed DyGAM-NS model optimizes the number of partitions and knots selection for natural cubic spline fitting, which allows for better capturing of the non-uniform and nonlinear biological characteristics of dimorphic transition. Moreover, our model optimizes gene-specific parameters, enabling the distinction of genes with different expression patterns during the dimorphic transition. Overall, the DyGAM-NS model achieved a balance between accuracy and recall, with both exceeding 90%, and outperformed commonly used models in terms of F1-score. The efficient and accurate identification of DTIGs can facilitate the exclusion of noise, resulting in improved accuracy in downstream gene expression pattern clustering and regulatory network construction. The DyGAM-NS model, as a novel approach for identifying dynamic temporal gene regulation, also offers a new insight into unraveling mechanisms of dimorphic transition or other complex biological processes.

In this study, we utilized temporal transcriptomes to investigate the dimorphic transition of *T. marneffei*, focusing on its distinct gene expression patterns at different stages of dimorphic transition. During the early stage of M-to-Y transition, we observed a substantial number of rapid upregulation genes associated with environmental adaptation, such as *pepA*, *ftrA*, *ftrC*, *sidA*, and *hmgR* (23, 24, 26). This observation suggests that *T. marneffei* undergoes a rapid adaptive response to the harsh environment within host macrophages. In contrast, we also identified genes with delayed response patterns, which are linked to characteristics specific to mycelial or yeast phase growth but unrelated to environmental stimuli. For instance, the gene *hgrA*, involved in cell wall regulation, remained unchanged until 48 hours into the Y-to-M transition. Functional analysis revealed that although *hgrA* is crucial for maintaining hyphal cell wall integrity, it does not directly participate in stress response (34), providing valuable insights into its upregulation during the later stage of the Y-to-M transition. Furthermore, we constructed a gene regulatory network for the dimorphic transition of *T. marneffei* using the DyGAM-NS model to identify key transcription factors. Interestingly, two zinc finger-containing transcription factors were found to potentially regulate the M-to- Y and Y-to-M transitions, respectively. These findings present new avenues for future research aimed at unraveling the regulatory mechanisms underlying the dimorphic transition of *T. marneffei*.

## Materials and Methods

### Strains media, and culture conditions

*Talaromyces marneffei* strain PM1, isolated from a talaromycosis patient without HIV infection in Hong Kong (8), served as the basis for our research. Previously, we generated *madsA* overexpression and knockout mutant strains (8, 41) derived from the wild-type PM1. All strains were maintained at a temperature of 25°C on Sabouraud Dextrose Agar (SDA) (Becton, Dickinson and Company, USA), which was also utilized for preparing conidial inocula. Sabouraud Dextrose Broth (SDB) (Becton, Dickinson and Company, USA) was employed for liquid cultures of the wild-type strain. To ensure long-term storage, mycelia from each strain were suspended in 25% (w/v) sterile glycerol and subsequently frozen at -80°C.

### Whole-genome sequencing of *T. marneffei* PM1

Initially, a conidia suspension of PM1, containing approximately 1 × 10^6^ conidia, was introduced into a 50 ml SDB culture. This culture was incubated at 37°C with shaking at 200 rpm for 7 days. Subsequently, 5 ml of the culture was transferred to a fresh 45 ml SDB and incubated overnight at 37°C. The resulting yeast cells were harvested through centrifugation at 3000 × g and washed twice with 1 × phosphate-buffered saline (PBS). Genomic DNA extraction was carried out using the Epicentre MasterPure Yeast DNA Purification Kit (Cat. MPY80020) according to the manufacturer instructions, with a minor modification: RNA was digested using RNase A, RNase T1, and RNase I instead of solely RNase A. The quantity and quality of the extracted genomic DNA were evaluated using the Qubit 4.0 fluorometer (Thermo Scientific, USA) and NanoDrop spectrophotometer (Thermo Scientific, USA). Whole genome sequencing was performed by utilizing the Illumina HiSeq X platform (350 bp insert size with 150 bp paired-end) and the PacBio Sequel platform (10 kb inserts library).

### *De novo* genome assembly and evaluation of quality

Subreads were obtained from the PacBio sequencing data using the SMRT analysis pipeline v5.1.0 (43). The Canu v1.6 algorithm (44) was utilized to generate a draft genome through an overlapping layout consensus (OLC) assembly approach, employing PacBio subreads. In this process, a genome size of 30 Mb was specified, and the longest 40× of subreads were selected as seed reads to generate error-corrected long reads. The draft assembly underwent further processing, including the filtration of bubble sequences and the identification of the mitochondria sequence by comparing it to the known *T. marneffei* sequence using BLASTN v2.12.0+ (45). To enhance the accuracy of the assembly, Arrow v2.0.2 (43) was employed based on PacBio reads, while Pilon v1.24 (46) was used based on Illumina reads for additional polishing. The completeness of the assembly was assessed by performing BUSCO v4 analysis (47) against the Eurotiales database.

### Temporal RNA preparation and sequencing

In the initial phase of the experiment, conidia of strain PM1 were inoculated onto SDA plates and cultivated at two different temperatures, 25°C and 37°C, for a duration of one week. Subsequently, the SDA plates incubated at 25°C were transferred to 37°C, while those cultivated at 37°C were transferred to 25°C. This process facilitated the growth of colonies representing the M-to-Y or Y-to-M transition growth forms, respectively. Total RNA samples were extracted at specific time points after the temperature switch, namely at 0, 1, 3, 6, 9, 12, 24, 36, 48, and 72 hours. In the case of the *madsA* overexpression and knockout strains, total RNA samples were extracted at 0 and 48 hours for the M-to-Y transition, and at 0 and 6 hours for the Y-to-M transition. For each time point, three independent biological replicates were used to extract total RNA using the E.Z.N.A. fungal RNA kit (Omega Bio-Tek). The quantification of RNA was performed using a Qubit 4.0 fluorometer (Thermo Scientific, USA). A total of 84 strand-specific libraries, with RNA integrity numbers (RIN) exceeding 6.5, were constructed and sequenced on an Illumina HiSeq X platform, resulting in approximately 20 million 150-bp paired-end reads for each sample. Prior to analysis, the reads were trimmed using Trimmomatic v0.38 (48) based on quality assessment carried out by FastQC v0.11.5 (https://www.bioinformatics.babraham.ac.uk/projects/fastqc/).

### Annotation of repeat sequences and protein-coding genes

Repeat regions in the assembly were masked using RepeatMaker v4.0.7 (http://repeatmasker.org) with a *T. marneffei*-specific repeat library generated by RepeatModeler v4.0.7 (http://repeatmasker.org/RepeatModeler/). Gene annotations were conducted using three distinct types of evidence. Firstly, the assembly was aligned to a protein database that combined the UniProt/Swiss-Prot protein database and all sequences of *Talaromyces marneffei* from the NCBI protein database, using Exonrate v2.2.0 (49). Additionally, temporal RNA-seq reads of *T. marneffei* PM1 from various time points during the dimorphic transition were *de novo* assembled with Trinity v2.3.2 (50). The resulting Trinity assembly was then passed through the PASA v2.5.2 pipeline (51) to generate transcript evidence. Furthermore, *ab initio* gene predictions were carried out using FGENESH (52) and BRAKER2 v2.1.6 (53). For FGENESH, the genome-specific parameters of *Penicillium* were employed, while BRAKER2 utilized RNA-seq reads to enhance the accuracy of gene predictions. The EVidenceModeler (EVM) v1.1.1 (54) was employed to integrate the three types of evidence into consensus gene structures, assigning a weight of 10 to transcript alignments, 5 to protein alignments, and 1 to *ab initio* gene predictions. The final gene sets were subjected to a BLASTP v2.12.0+ (45) search against the NCBI non-redundant (NR) protein database. The domains of these genes were annotated using InterProScan v5.56-89.0 (55), which utilizes publicly available databases. Functional annotations were then performed using BLAST2GO v5.2.5 (56), incorporating the results obtained from both BLASTP and InterProScan analyses.

### RNA-seq data analysis

RNA-seq short reads were aligned to the annotated genomes using STAR v2.7.9a (57) to obtain read mappings. The read count for each gene was calculated using FeatureCount v2.0.3 (58), specifying a library by using “-s 2”. The expression level of genes was quantified using TPM (transcripts per million). Genes with a mean TPM greater than one at least once during the dimorphic transition were selected for subsequent analysis. The raw counts were then normalized using the median of ratios method implemented in DESeq2 (59). For the time-course RNA-seq data, principal component analysis (PCA) was performed using the prcomp function in R v4.2.2. Visualization of the PCA results was achieved using the ggplot2 package. The heatmap depicting gene expression levels during the dimorphic transition was generated using the pheatmap package. In the case of RNA-seq data from *madsA* overexpression and knockout strains, differentially expressed genes were identified using the DESeq2 method, applying a false discovery rate (FDR) threshold of less than 0.05 and a fold change threshold greater than 2.

### Identification of dimorphic transition induced genes (DTIGs)

We utilized a dynamic optimized generalized additive model with natural cubic smoothing (DyGAM-NS) approach to capture the nonlinear relationship between gene expression profiles and time during the dimorphic transition (**Eq. 1**).

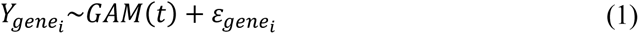

To account for the discrete nature and overdispersion of read counts, we employed a negative binomial (NB) distribution with gene-specific means and dispersion, estimated using DESeq2 (**Eq. 2**).

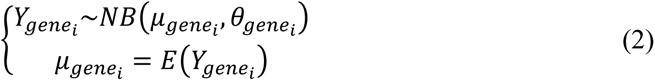

The flexibility of the DyGAM-NS method allowed us to adjust the number and placement of knots (**Eq. 3**). To determine the optimal parameters for each gene, we evaluated models with all possible combinations of knots using the corrected Akaike information criterion (AICc).

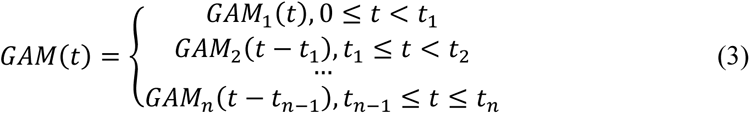

The DyGAM-NS model was fitted using the gam function in the mgcv package (60), and AICc values were calculated using the AICc function in the AICcmodavg package. Finally, we filtered the dimorphic transition induced genes (DTIGs) based on criteria of FDR less than 0.05, deviance explained greater than 0.75, and fold change greater than 2.

### Evaluation and comparison of the DyGAM-NS model

To evaluate the precision of the DyGAM-NS model, we constructed a gene set by identifying differentially expressed genes through pairwise comparisons using DESeq2. For the evaluation of recall, we identified high variable genes based on the methodology described in a previous study (18). In addition to the DyGAM-NS model, we employed seven other time-course analysis tools in this study, all implemented in R with their default parameters. Here is a brief overview of the methods used: impulse model using ImpulseDE2 (20), empirical Bayes autoregressive hidden Markov model using EBSeqHMM (12), polynomial regressive model using Next maSigPro (10), autoregressive model using stat, cubic spline model using edgeR (61) and splines, generalized linear model using edgeR, and linear model using limma (62).

### Clustering gene expression patterns of DTIGs

To analyze the expression pattern of DTIGs, we employed the trimmed *k*-means method from the tclust package (22). This method utilized the fitted values of the DyGAM-NS model for each gene, where the fitted values were standardized to have a zero mean and unit variance across samples for each gene. The optimal number of clusters was determined based on the within sum of squares.

### Construction of gene regulatory networks

Gene regulatory networks were constructed using GRNboost2 v0.1.6 (38) based on the time-course expression data sets. The list of transcription factors (TFs) used as input for GRNboost2 was identified through the InterPro database (IPR) and compiled from previous studies (39). The final gene regulatory network was obtained by retaining edges with the top 5% cumulative weight. The betweenness centrality of each gene was calculated using the igraph package in R. Visualization of the gene regulatory network was performed using the NetworkD3 package in R. The motifs associated with each regulatory gene in the network were identified using MEME (63), utilizing the upstream 500-bp regions of the corresponding target genes.

### Data availability

The whole genome sequencing data have been deposited in the Genome Sequence Archive in National Genomics Data Center, China National Center for Bioinformation under the accession PRJCA002718. The genome assembly has been deposited at GenBank database under the accession JPOX02000000. The time-course RNA-seq data are available from the National Center for Biotechnology Information BioProject database under the accession PRJNA970557.

## Acknowledgements

This work was funded by National Natural Science Foundation of China (31970008, Ence Yang).

